# Search for top-down and bottom-up drivers of latitudinal trends in insect herbivory in oak trees in Europe

**DOI:** 10.1101/2020.02.25.964213

**Authors:** Elena Valdés-Correcher, Xoaquín Moreira, Laurent Augusto, Luc Barbaro, Christophe Bouget, Olivier Bouriaud, Manuela Branco, Giada Centenaro, György Csóka, Thomas Damestoy, Jovan Dobrosavljević, Mihai-Leonard Duduman, Anne-Maïmiti Dulaurent, Csaba B. Eötvös, Maria Faticov, Marco Ferrante, Ágnes Fürjes-Mikó, Andrea Galmán, Martin M. Gossner, Arndt Hampe, Deborah Harvey, Andrew Gordon Howe, Yasmine Kadiri, Michèle Kaennel-Dobbertin, Julia Koricheva, Alexander Kozel, Mikhail V. Kozlov, Gábor L. Löveï, Daniela Lupaştean, Slobodan Milanović, Anna Mrazova, Lars Opgennoorth, Juha-Matti Pitkänen, Anna Popova, Marija Popović, Andreas Prinzing, Valentin Queloz, Tomas Roslin, Aurélien Sallé, Katerina Sam, Michael Scherer-Lorenzen, Andreas Schuldt, Andrey Selikhovkin, Lassi Suominen, Ayco J. M. Tack, Marketa Tahadlova, Rebecca Thomas, Bastien Castagneyrol

## Abstract

**Aim:** The strength of species interactions is traditionally expected to become stronger toward the Equator. However, recent studies have reported opposite or inconsistent latitudinal trends in the bottom-up (plant quality) and top-down (natural enemies) forces driving insect herbivory, possibly because these forces have rarely been studied concomitantly. This makes previous attempts to understand the effect of large scale climatic gradients on insect herbivory unsuccessful.

**Location:** Europe

**Time period:** 2018-2019

**Major taxa studied:** *Quercus robur*

**Methods:** We used scholar-based citizen science to simultaneously test for latitudinal variation in plant-herbivore-natural enemy interactions. We further investigated the underlying climatic factors associated with variation in herbivory, leaf chemistry and attack rates in *Quercus robur* across its complete latitudinal range in Europe. We quantified insect herbivory and the occurrence of specialist herbivores as well as leaf chemistry and bird attack rates on dummy caterpillars on 261 oak trees.

**Results:** Climatic factors rather than latitude *per se* were the best predictors of the large-scale (geographical) variation in the incidence of gall-inducers and leaf-miners as well as of leaf nutritional quality. However, insect herbivory, plant chemical defences (leaf phenolics) and bird attack rates were not influenced by latitude or climatic factors. The incidence of leaf-miners increased with increasing concentrations of hydrolysable tannins and decreased with those of condensed tannins, whereas the incidence of gall-inducers increased with increasing leaf soluble sugar concentration and decreased with increasing leaf C:N ratios. However, neither other traits nor bird attack rates varied with insect herbivory.

**Main conclusions:** These findings help to refine our understanding of the bottom-up and top-down mechanisms driving geographical variation in plant-herbivore interactions, and urge for further examination of the drivers of insect herbivory on trees.

## Introduction

Ecological theory predicts that the strength of species interactions increases toward the Equator due to warmer temperatures, longer growing seasons, and higher species abundance and diversity at lower latitudes (Janzen, 1970; Schemske *et al*., 2009). Plant species at lower latitudes commonly experience higher rates of herbivory than plants growing further away from equator (Coley & Barone, 1996; Schemske *et al*., 2009; Lim *et al*., 2015; Moreira *et al*., 2018) and thus tropical plant species may evolve higher levels of anti-herbivore defences (Johnson & Rasmann, 2011; Pearse & Hipp, 2012; Abdala-Roberts *et al*., 2016). While early experimental studies reported patterns supporting these predictions (Coley & Aide, 1991; Coley & Barone, 1996; Dyer & Coley, 2009), several studies over the last decade have found either no evidence for a latitudinal gradient in herbivory and plant defences (Moles *et al*., 2011) or increase in herbivory and defences with latitude (Woods *et al*., 2012; Moreira *et al*., 2018, 2020). Given these inconsistencies, it is of great importance to identify the mechanisms underlying the substantial variation in herbivory and plant defences across latitudes, as insect herbivory is an important ecological process that affects primary productivity by altering the recruitment, mortality and growth of plants.

Latitudinal gradients can be used as ‘natural laboratories’ to study the relationship between climate and plant-herbivore interactions (De Frenne *et al*., 2013; Kozlov *et al*., 2015; Lim *et al*.,2015; Moreira *et al*., 2018). In the northern extratropical hemisphere, mean annual temperature drops by 0.73 °C and mean annual precipitation by 4.04 mm per degree of latitude northward (De Frenne *et al*., 2013). Latitudinal variation in plant-herbivore interactions is therefore generally associated with large-scale variability in climatic conditions (Moreira *et al*., 2018) and numerous studies demonstrate an effect of temperature and precipitation on plant traits (e.g. leaf N, phenolic compounds) (Chen *et al*., 2013; Holopainen *et al*., 2018; Gely *et al*., 2019) and herbivory (Jamieson *et al*., 2015; Gely *et al*., 2019). However, many regions deviate from the global trend in temperature and precipitation toward higher latitudes due to their proximity to oceans or the presence of mountains (De Frenne *et al*., 2013), which can markedly change the relationship between latitude and plant-herbivore-predator interactions (Roslin *et al*., 2017; Loughnan & Williams, 2019; Moreira *et al*., 2019). Thus, further studies should not only rely on latitudinal clines to infer the effect of climate on plant-herbivore interactions, but should also stretch latitudinal gradients longitudinally to better capture the diversity of climatic conditions in which plant-herbivore interactions are embedded and use climatic factors such as temperature and precipitation rather than latitude *per se* as predictors (Anstett *et al*., 2016).

Recent work identified several potential sources of variation in the direction and strength of latitudinal gradients in herbivory and plant defences (Johnson & Rasmann, 2011; Anstett *et al*.,2016). First, theory on latitudinal gradients in herbivory and plant defences assumes a plant-centred equilibrium in which plants at low latitudes have adapted to higher herbivory levels by evolving greater defences. However, most studies have measured either herbivory patterns or plant defences, but not both (but see Anstett *et al*., 2015; Moreira *et al*., 2018), leading to an incomplete understanding of the relationship between latitudinal clines and plant-herbivore interactions. Second, little attention has been paid to latitudinal variation in tritrophic dynamics (Roslin *et al*., 2017). Herbivore natural enemies, however, can drastically modify tritrophic interactions by suppressing herbivore populations or reducing herbivore feeding (Rosenheim, 1998; Maguire *et al*., 2015). In the few published studies exploring latitudinal patterns in natural enemy activity, authors have found no variation in parasitism (Dyer & Coley, 2002; Moreira *et al*., 2015), lower attack rates pressure by ants (Roslin *et al*., 2017), and higher (Zvereva *et al*., 2019) or no variation (Roslin *et al*., 2017) in attack rates by birds with increasing latitude. Thus, considering bottom-up and top-down forces simultaneously is crucial for a complete understanding of latitudinal clines in plant-herbivore interactions.

We aimed to test for latitudinal variation in plant-herbivore-natural enemy (i.e. tritrophic) interactions, as well as the underlying climatic factors associated with variation in herbivory and defences in the pedunculate oak (*Quercus robur*), a long-lived, common European tree. In particular, we asked the following questions: (1) Are there latitudinal clines in herbivory? (2) Is latitudinal variation in leaf chemical traits (bottom-up effects) and/or herbivore attack rates (topdown effects) associated with latitudinal variation in herbivory? (3) Are climatic correlates of latitude associated with clines in herbivory, leaf chemical traits and attack rates pressure? We used data collected by professional scientists and schoolchildren across major parts of the geographical distribution range of *Q. robur*. We quantified insect leaf damage, leaf chemical traits (phenolics, soluble sugars and nutrients) and attack rates on dummy caterpillars placed on mature oak trees. Overall, our study aims to identify latitudinal patterns in plant-herbivore-predator interactions and helps to refine our understanding of bottom-up and top-down mechanisms that may drive geographical variation in plant-herbivore interactions.

## Material and methods

The present study involved 30 professional scientists from 14 countries and 82 school teachers (with their pupils) from 10 countries in 2018 and 2019, giving a total of 112 partners from 17 European countries, covering most of the native geographic range of the pedunculate oak **(Figure 1)**. Every partner received detailed instructions at the beginning of the project (Castagneyrol *et al*., 2019). Here, we only provide a summary of these instructions. Only project partners who provided data that could be used in the present article were considered.

**Figure 1.**
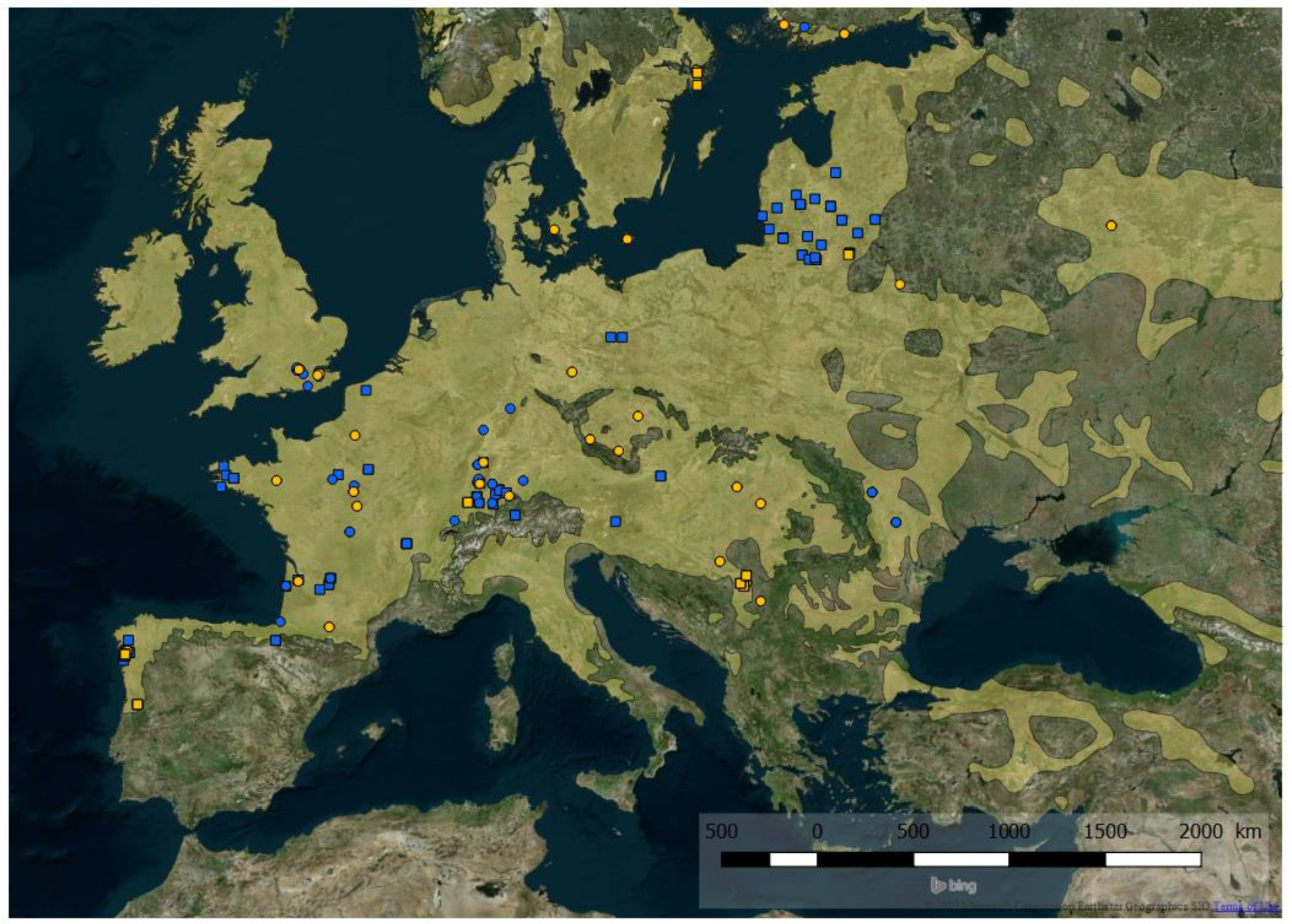
Distribution range of *Quercus robur* L. (shaded in yellow) and locations of trees sampled by professional scientists (orange symbols) and schoolchildren (blue symbols) in 2018 (circles) and 2019 (squares). Additional maps showing oak trees used for estimating leaf herbivory, attack rates on dummy caterpillars and trait analyses are provided in supplementary material (Figure S1).

### Target species

The pedunculate oak, is one of the dominant deciduous tree species in European forests with high economic and symbolic value (Eaton *et al*., 2016). Its distribution ranges from Central Spain (39°N) to southern Fennoscandia (62°N), thus experiencing variable climatic conditions(Petit *et al*., 2002). This species supports a large community of specialist and generalist herbivore insects; especially suckers, chewers, skeletonizers, gall-inducers and leaf-miners that are mainly active between the time of leaf burst and fall (Southwood *et al*., 2005; Moreira *et al*., 2018), as well as xylophagous species (Marković & Stojanović, 2011). The wide distribution of pedunculate oak and the high diversity of associated herbivorous insects makes it a suitable model species for research on the effect of climate on biotic interactions.

In total, the study included 261 mature oak trees surveyed by professional scientists (n = 115) and schoolchildren (n = 146) in 2018 (n = 149) and 2019 (n = 113) (**Figure 1**). However, not every partner measured or provided material allowing measuring herbivory, bird attack rates and leaf chemistry simultaneously on every tree (Figure S1, supplementary material).

### Attack rates on dummy caterpillars

To control for latitudinal variation in environmental conditions, we matched the start of the experiment in each locality to the phenology of the local oak trees. Six weeks after oak budburst, partners installed 20 dummy caterpillars per tree, *i.e*., five caterpillars on each of four branches (facing north, south, east and west) and a minimum distance of 15 cm between caterpillars. We also verified that the starting date of budburst and the latitude were positively correlated (Pearson *r* = 0.45, *P* < 0.05).

In orther ot measure bird attack rate, the project coordinators provided the same green plasticine (Staedler, Noris Club 8421, green[5]) to all partners to make the caterpillars. In order to standardize caterpillar size among partners, we made caterpillars from a 1 cm diam ball of plasticine, and gently pressed/rolled this along a 12 cm long metallic wire until a 3 cm long caterpillar was obtained, with the wire in its center. Partners attached the caterpillars to branches by twisting the wire and left the caterpillars on trees for 15 days before recording predation marks. A second survey using the same procedure immediately followed the first one. In 2018, schoolchildren photographed every caterpillar with the suspected attack marks from any potential predatory taxon. In 2019, both schoolchildren and professional scientists sent caterpillars back to the project coordinators.

In order to be consistent and reduce bias due to multiple observers, photos and dummy caterpillars were screened by a single trained observer (first author, EVC). For each oak tree and survey period, we assessed attack rate as the proportion of dummy caterpillars with at least one attack mark. Although we asked partners to record attack rate marks left by different types of predators (in particular birds and arthropods), attacks by arthropod predators could not be verified on photos because of their low resolution. In addition, the relevance of marks left by arthropods on plasticine model prey has recently been questioned, in particular after mandibular marks were observed on lizards or frog models (Rößler *et al*., 2018). For these reasons, we decided to discard arthropod attack rate from the study and focused on marks that were unambiguously attributed to birds, *i.e*., conic holes or V-shaped beak marks. Attack marks left by reptiles or rodents were also disregarded, because only a few caterpillars were attacked by these potential predators. Most bird marks were directed towards the head or the body centre of the dummy caterpillars, which is typical to bird attacks and indicates prey recognition(Rößler *et al*., 2018). We therefore refer to the proportion of dummy caterpillars with such marks as bird attack rate.

Between 2018 and 2019, 137 partners installed 12,760 dummy caterpillars on 319 oak trees. Despite clear instructions regarding caterpillar installation, removal and conditioning prior to shipping, the material sent by 22 school partners was of poor quality (with no particular geographic bias) such that only caterpillars returned by 115 partners (*i.e*., 78.4%, collected on 254 oak trees) were screened for attack marks and included in subsequent analyses (**Table S1; Figure 1**).

### Insect herbivory

Professional scientists and schoolchildren were instructed to collect oak leaves after the second bird attack rate survey, *i.e*., roughly 10 weeks after oak budburst, on the same branches where dummy caterpillars were installed. They haphazardly collected 30 leaves per branch, totalling 120 leaves from which they blindly drew 60 leaves. Professional scientists oven-dried leaves for a minimum of 48 h at 45°C immediately after collection, and leaves collected by schoolchildren were oven dried upon receipt by the project coordinators, to ensure optimal conservation prior herbivory assessment.

For each leaf, we visually assessed insect herbivory as the percentage of leaf area removed by leaf chewers and skeletonizers following eight levels of defoliation (0%, 0.1-5%, 5.1-10%, 10.1-15%, 15.1-25%, 25.1-50%, 50.1-75%, and >75.1%). Herbivory assessment was made by two trained observers who were blind to leaf origin to reduce unconscious bias. We then averaged herbivory at tree level using the midpoint of each percentage class to obtain a mean value per tree. This measurement also included the surface covered by leaf mines, but we did not consider punctures made by sap feeders. Additionally, we also scored the presence of gall-inducers and leaf-miners at leaf level and calculated the incidence of gall-inducers and leaf-miners as the proportion of leaves with mines or galls.

### Leaf chemical traits

We used leaves sent by professional scientists and schoolchildren in 2018 to quantify several leaf chemical traits typically recognized as deterrents against insect herbivores for several oak species. Details of procedures used to analyse chemical leaf traits are reported in online supplementary material S1.

We quantified leaf phenolics as toxic defensive metabolites (Moreira *et al*., 2018). We used only leaves collected by professional scientists in 2018. From each tree, we selected 10 mature, dried leaves with no evidence of insect damage and grounded them to fine powder. We identified four groups of phenolic compounds: flavonoids, ellagitannins and gallic acid derivates (“hydrolysable tannins” hereafter), proanthocyanidins (“condensed tannins” hereafter) and hydroxycinnamic acid precursors to lignins (“lignins” hereafter) (see Supplementary Material S1 for further details).

We quantified C:N ratio, N:P ratio, cellulose and soluble sugars as proxies for leaf nutritional quality to herbivores (Moreira *et al*., 2019). We measured these traits on leaves collected by both professional scientists and schoolchildren. We grounded the 60 oven dried leaves on which we scored herbivory to fine powder such that leaf nutritional traits reflected the content of leaves with different amounts of herbivore damage (see Supplementary Material S1 for further details).

### Statistical analysis

We were primarily interested in testing the interactive effects of climate and leaf chemistry on herbivory and bird attack rate. Thus, we focused on temperature and precipitation that we obtained from the WorldClim database (Hijmans *et al*., 2005) based on oak coordinates as retrieved on Google maps by project partners, so that the sampled geographic gradients were taken as proxies for climatic gradients. Specifically, we extracted the mean temperature and precipitation from April to June, which roughly corresponded to the period when caterpillars were present on trees, irrespective of latitudinal cline in moth phenology. Yet, because latitude can have interactive effects on temperature and precipitation, we tested the effect of geography and climate separately.

We were interested in whether bird attack rate or leaf chemistry mediated the effect of climate on insect herbivory. Yet, leaf chemistry were only measured on a subset of trees such that we could not link herbivory with its top-down and bottom-up drivers using the complete dataset. Therefore, we built three types of Linear Mixed-effects Models (LMM): (i) a geographic model analysing the effect of latitude on herbivory, leaf chemistry and bird attack rate, (ii) a climatic model in which we substituted latitude with climatic data (temperature and precipitation) and (iii) an abiotic and biotic model analysing the effects of leaf chemistry and bird attack rate together with temperature and precipitation or latitude (both linear and quadratic) on herbivory.

In every LMM, we used Partner ID as a random factor to account for the fact that some partners surveyed multiple trees. For instance, the geographic models were of the form:

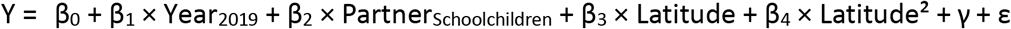

where Y was the response variable, β_i_ model coefficient parameter estimates, Partner_Schoolchildren_ was the effect of partner type (the estimate for schoolchildren being compared with the estimate for professional scientists that was included in the intercept), Year_2019_ was the effect of each year (2018 contrasted with 2019), Latitude (and their quadratic terms) the geographic conditions around sampled oak trees, σ^2^_Partner ID_ the random effect of Partner ID (assuming that γ ∈ N(0, σ^2^_Partner ID_) and ε the residuals (assuming ε∈ N(0, σ^2^_e_)). When Y was bird attack rate, we added the survey (first vs. second) as a fixed effect and Tree ID as a random effect nested within Partner ID to account for repeated measurements on the same trees. Partner type was added to adjust for differences between the two partner types. When needed, we used arcsine square-root (bird attack rate) or logarithmic (insect herbivory, soluble sugars, N:P ratio and leaf defences) transformations of the response variable to satisfy model assumptions.

We ran geographic and climatic models on the complete dataset including 2018 and 2019 data collected by both professional scientists and schoolchildren. Note that because not every partner provided reliable data on both bird attack rates and herbivory, the sample sizes differed between models using bird attack rate or herbivory as response variables **(Figure 1, Figure S1)**. We ran the geographic and climatic models on leaf phenolics as well as the biotic model on the 2018 data collected by scientific partners only, as we did not quantify leaf defences on leaves collected and sent by schoolchildren.

The tree-level response variables for each year and survey period (Y) were either herbivory (% leaf area removed by herbivores), the presence of leaf-miners or gall-inducers (proportions), mean bird attack rate (ratio of % attacked caterpillars on exposition period) or leaf chemistry (C:N ratio, N:P ratio, soluble sugar content [g L^-1^], cellulose content (g), concentrations of condensed or hydrolysable tannins, flavonoids or lignins [mg g^-1^ d.m.]). We scaled and centred every continuous predictor prior to modelling to facilitate comparisons of their effect sizes. We used LMM with a Gaussian error distribution, with the exceptions of geographic, climatic and process-based models with the incidence of leaf-miners or gall-inducers as response variables. In these cases, we used Generalized LMM with a binomial error distribution.

We analysed the data within the information theory framework (Burnham & Anderson, 2002). We first built a set of models including geographic and climatic models as well as nested models for each response variable separately. Biotic models were run on the subset of samples where all data were measured simultaneously. We then applied a procedure of model selection based on AIC criteria corrected for small sample size (AICc). In the first step, we ranked the models according to the difference in AICc between a given model and the model with the lowest AICc (ΔAICc). Models within 2 ΔAICc units of the best model (*i.e*., the model with the lowest AICc) are generally considered as equally likely. We also computed AIC weight (w_i_) that is the probability a given model to be the best model, as well as the relative variable importance RVI as the sum of w_i_ of every model including this variable. When several models compete with the best model (*i.e*., when multiple models are such that their ΔAICc < 2), we applied a procedure of multimodel inference building a consensus model including the variables in the set of best models. We then averaged their effect size across all the models in the set of best models, using variable w_i_ as a weighting parameter (*i.e*., model averaging). We considered that a given predictor had a statistically significant effect on the response variable when its confidence interval excluded zero.

In the results section, we report model AICc, ΔAICc and *w_i_* for every model, as well as averaged coefficient parameter estimates and variable importance for all variables present in the set of competing models. When appropriate, we plotted the relationship between raw data and explanatory variables together with the predictions of simplified models, holding undisplayed predictors constant. All analyses were run in the R language environment (R Core Team, 2018) with packages *MuMIn* (Barton, 2018) and *lme4* (Bates *et al*., 2018).

## Results

### Latitudinal and climatic gradients in insect herbivory, leaf chemistry and bird attack rates

Insect herbivores damaged on average (± se) 8.7 ± 0.4 % of leaf area (n = 182 trees, see **Table S1** for details). Model simplification identified the null model as the best model given the model set, indicating that none of predictors had a consistent effect on insect herbivory (**Table S2**). Insect galls were present on 7.1 ± 0.6 % of inspected leaves (n = 182, **Table S1)**. In the set of best models **(Table S2)**, gall-inducers increased non-linearly with increasing spring temperature, with a steeper slope at higher temperatures **(Figure 2A)**. Gall-inducers peaked at intermediate levels of spring precipitation **(Figure 2B)** and was on average higher in 2018 than in 2019 **(Figure S2)**. Other predictors had no significant effects on gall-inducers. Leaf-miners were present on 18.2 ± 1.3 % of inspected leaves (**Table S1**). In the set of best models (**Table S2**), leaf-miners peaked at intermediate mean spring temperatures (**Figure 2C**). Leaf-miners decreased non-linearly with increasing spring precipitation, with a steeper slope at lower precipitations (**Figure 2D**). Leaf-miners was significantly higher in 2018 than in 2019 (**Figure S2**) and higher in leaves sampled by professional scientists than in those sampled by schoolchildren.

**Figure 2.**
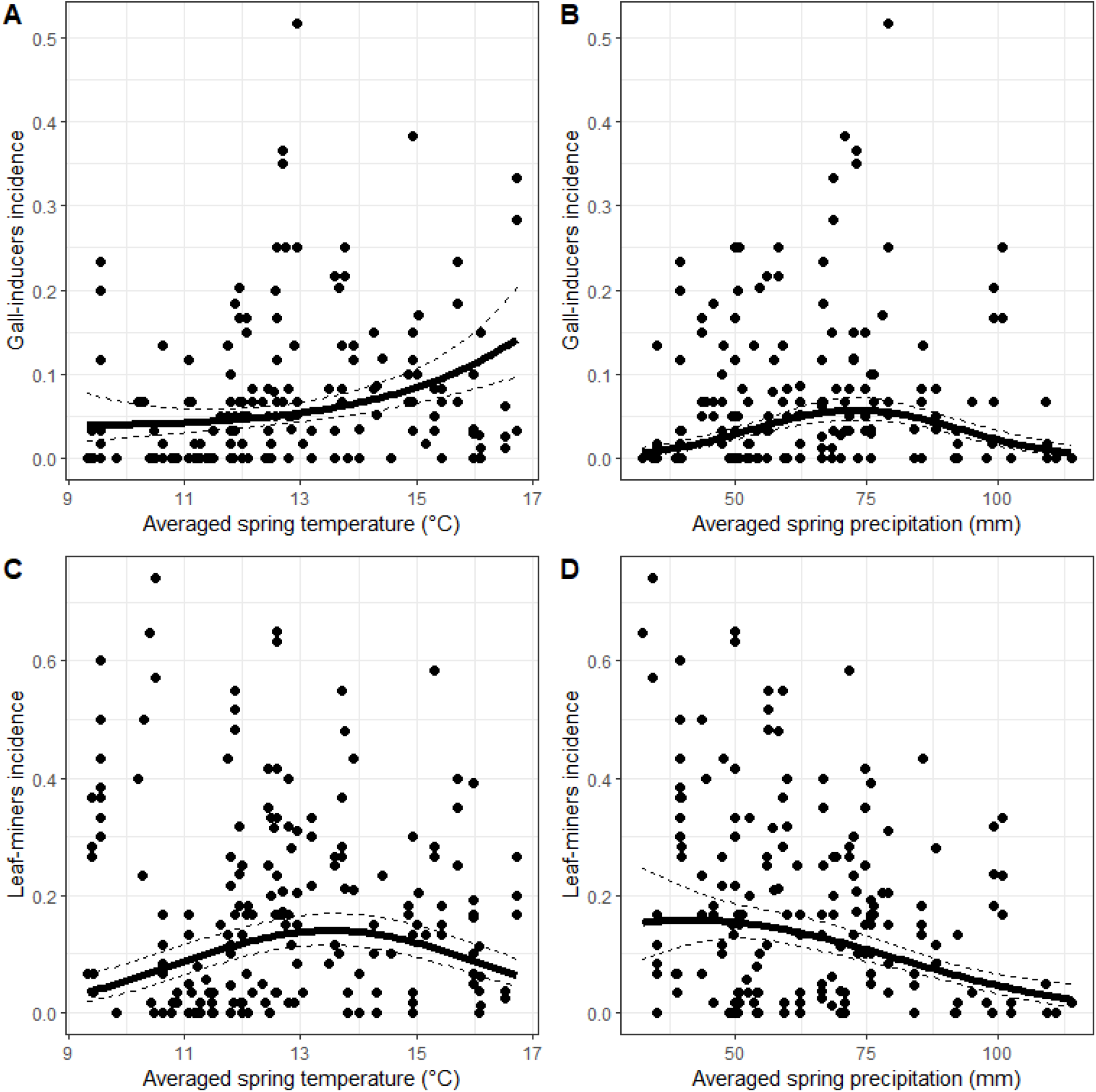
Effects of temperature and precipitation on gall-inducers (A, B) and leaf-miners (C, D) incidences. Dots represent raw data averaged at the tree level. Solid and dashed lines represent model predictions (and corresponding standard error) for temperature and precipitation calculated after other significant variables (see Table A5) were set to their mean value. Only statistically significant relationships are shown. Regression line equations are as follows: **A**, y = −2.28 + 0.34 · x + 0.05 · x^2^; **B**, y = −2.28 + 0.39 · x - 0.35 · x^2^; **C**, y = −1.36 + 0.23 · x - 0.29 · x^2^; **D**, y = −2.25 + 0.21 · x - 0.28 · x^2^.

Climate and latitude significantly affected some oak nutritional traits, but not phenolic compounds (**Table S1**). Specifically, leaf soluble sugar content (3.7 ± 0.2 g·L^-1^, n = 114, **Table S1**) decreased with increasing precipitation (**Figure 3A**). Leaf C:N rate (18.6 ± 0.2, n = 114, **Table S1**) increased non-linearly with latitude (with concave slope, **Figure 3B**) and was on average lower in leaves collected by professional scientists than those collected by schoolchildren. None of the predictors had a significant effect on N:P or cellulose content **(Table S1)**.

**Figure 3.**
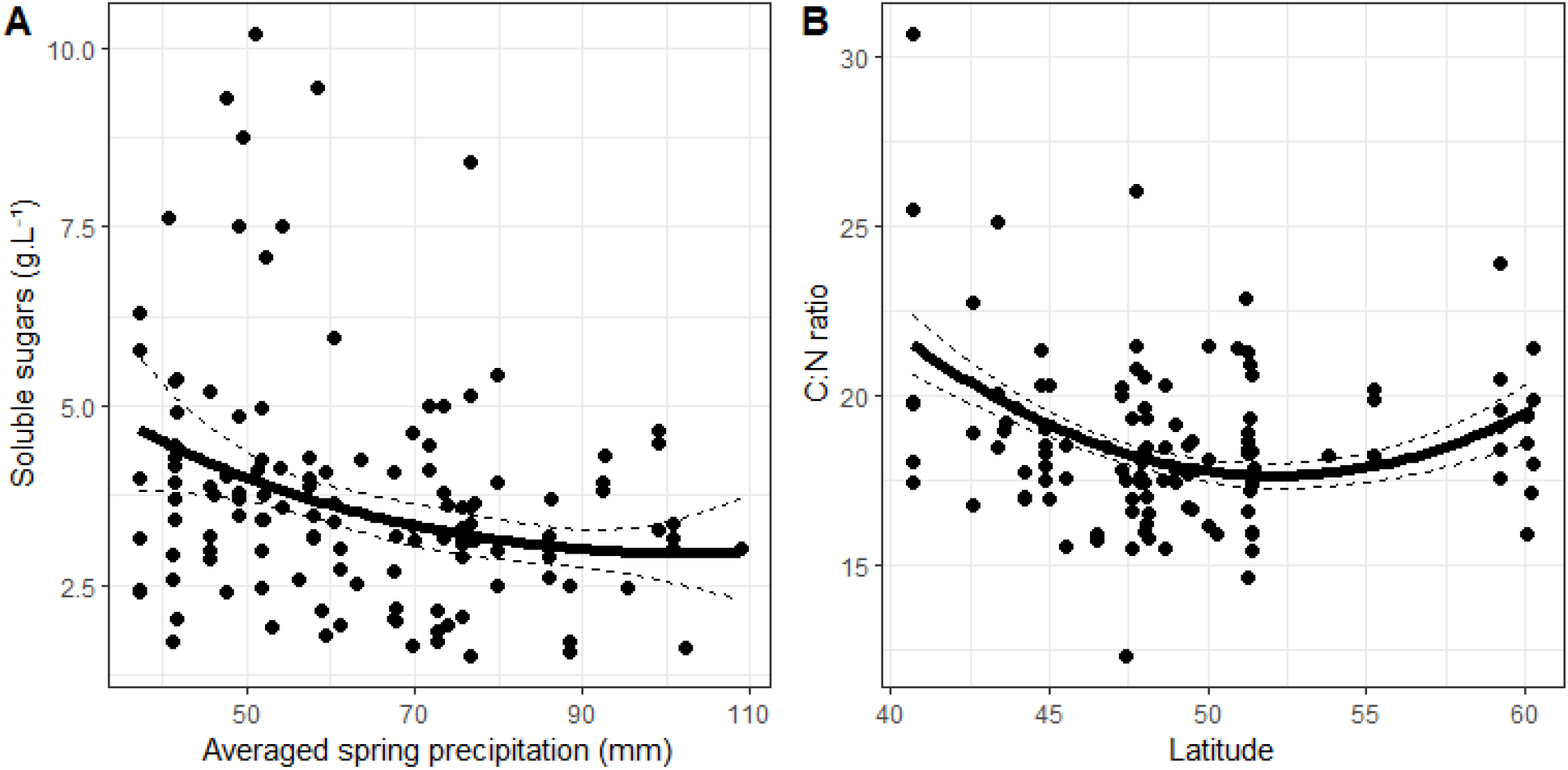
Effect of precipitation and latitude on soluble sugar (A) and C:N ratio (B) on leaves, respectively. Dots represent raw data averaged at tree level. Solid and dashed lines represent model predictions (and corresponding standard error) for temperature and precipitation calculated after other significant variables (see Table A6) were set to their mean value. Only significant relationships are shown. Regression line equations are as follows: **A**, y = 1.51 - 0.12 · x + 0.03 · x^2^; **B**, y = 1.52 - 0.03 · x + 0.03 · x^2^.

From a total of 10,000 dummy exposed caterpillars, 2,390 had bird marks (*i.e*., 23.9%). Model selection identified the null model as the best model, with no other competing model within two units of ΔAICc of the best model. This indicates that none of the predictors had a significant effect on bird attack rate on oaks at European scale.

### Mechanisms underlying latitudinal and climatic variation in herbivory

Model selection based on this data subset identified the null model as the best model, indicating that none of the examined predictors had a significant effect on insect herbivory **(Table S3)**.

When leaf chemistry were included in the model, gall-inducers incidence increased with increasing soluble sugar concentration and decreased with increasing C:N ratio **(Figure 4)**. The positive relationship between temperature and gall-inducers remained significant, suggesting independent effects of C:N ratio and temperature on gall-inducers. Gall-inducers incidence also increased with increasing latitude. The relative importance of leaf chemistry predictors (RVI > 0.8) was much higher than that of latitude or temperature (RVI < 0.4, **Figure S5**).

Leaf-miners incidence increased with increasing concentration of hydrolysable tannins and decreased with increasing concentration of condensed tannins. Other predictors had no significant effects on leaf-miners (**Figure 4**; **Table S3**).

**Figure 4.**
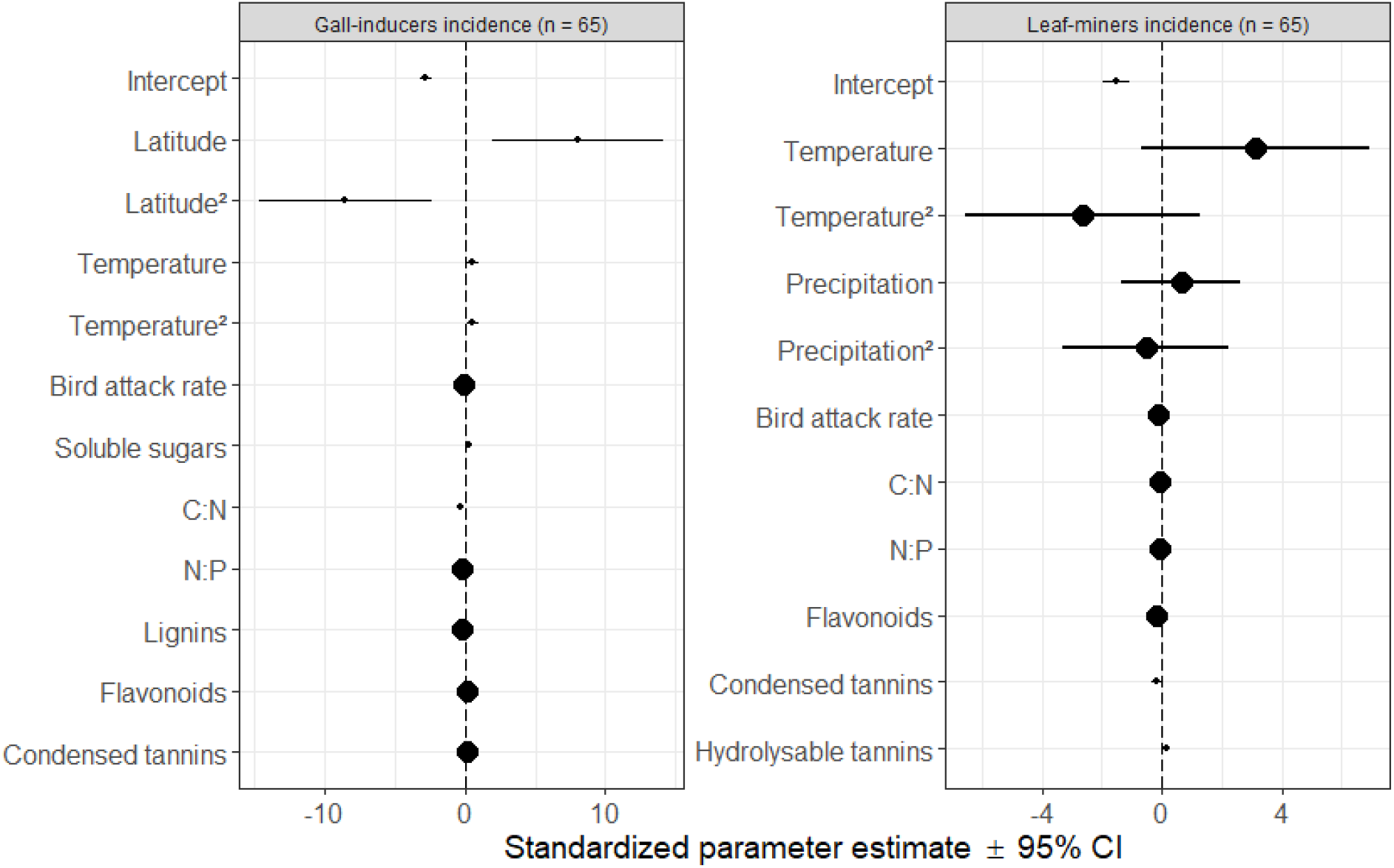
Effects of latitude, temperature, precipitation and the proportion of leaf chemistry on gall-inducers (left) and leaf-miners (right) incidences. Circles and error bars represent standardized parameter estimates and corresponding 95% CI. The vertical dashed line centred on zero represents the null hypothesis. Small and big circles represent significant and non-significant effect sizes, respectively.

## Discussion

### Latitudinal and climatic gradients in insect herbivory, plant chemical traits and bird attack rates

Variation in the incidence of gall-inducers and leaf-miners was associated with variation in temperature and precipitation, rather than with latitude *per se* (Anstett *et al*., 2018; Moreira *et al*., 2018; Loughnan & Williams, 2019). Climatic effects on herbivory were, however, contingent on herbivore feeding guild, with significant effects of climatic conditions detected for gall-inducers and leaf-miners, but not for leaf herbivory (leaf chewers and skeletonizers). The absence of variation in insect herbivory may be because herbivore guilds vary along abiotic conditions, so grouping different types of herbivores may avoid detecting patterns of each herbivore type (Abdala-Roberts *et al*., 2015; Moreira *et al*., 2015; Anstett *et al*., 2016) as each herbivore guild may respond differently to climatic clines (Anstett *et al*., 2016). The incidence of both gall-inducers and leaf-miners increased non-linearly with increasing temperature, but the shape of this relationship was accelerating (*i.e*., concave) for gall-inducers but decelerating (*i.e*., convex) for leaf-miners (**Figure 5**). Although we did not identify leaf-miners identity, this result is in line with that of Kozlov *et al*. (2013) who found that in the northern Europe, the diversity of leaf miners on birch trees increased linearly toward lower latitudes and that it was most likely associated with the direct impact of temperature, especially during cold years. At the same time, in our study the incidence of gall-inducers peaked at intermediate precipitation (Blanche & Ludwig, 2001; Leckey *et al*., 2014) whereas leaf-miners exhibited the opposite pattern. It has been hypothesized that gall-inducers and leaf-miners have evolved partly as adaptation to abiotic factors such as UV radiation and desiccation (Fernandes & Price, 1992; Connor *et al*., 1997; Danks, 2002). If so, gall-inducers and leaf-miners may be expected to be more common in the warmest and driest parts of pedunculate oak range and at low latitudes where the light intensity is markedly higher (Fernandes & Price, 1992; Lara & Fernandesrs, 1996; Price *et al*., 1998; Cuevas-Reyes *et al*., 2004). However, even within the gall-inducer and leaf-miner groups, relationships to climate are highly variable among species and years (Blanche, 2000; Sinclair & Hughes, 2010; Kozlov *et al*., 2013, 2016), thus suggesting that other factors are also important in the incidence of gall-inducers and leaf-miner herbivores.

**Figure 5.**
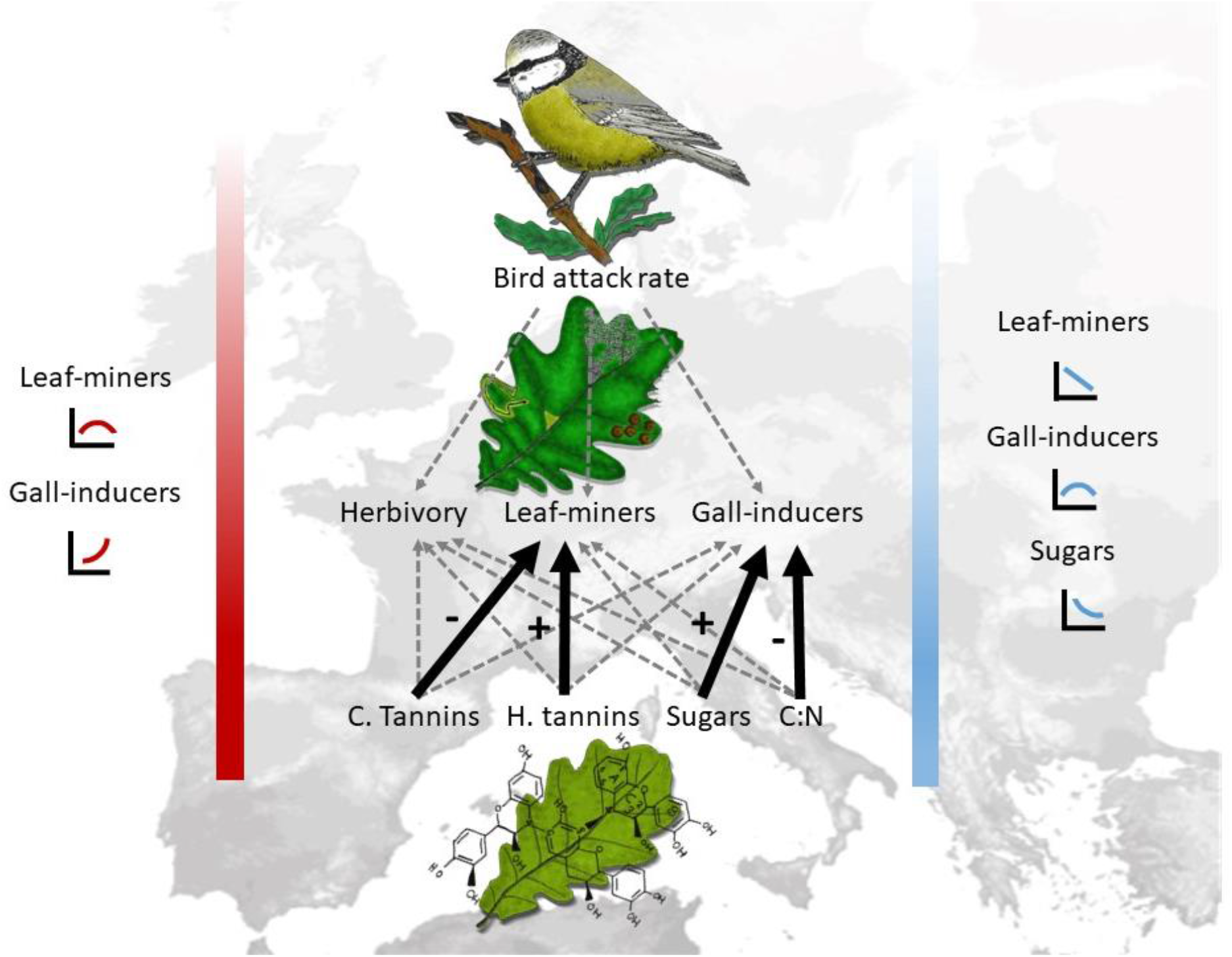
Summary illustrating plant-herbivore-predator relationships along a latitudinal gradient in Europe. The red and blue bands denote the variation in mean spring temperature and precipitation, respectively. The two figures on the left represent the correlation between the mean spring temperature and the incidence of gall-inducers and leaf-miners. The three figures on the right correspond with the correlation between mean spring precipitation and the incidence of leaf-miners and gall-inducers and the concentration of soluble sugar in leaves. Solid black arrows represent significant positive (+) or negative (-) relationships; dashed grey lines indicate non-significant relationships.

We did not find detectable latitudinal and climatic gradients in plant defences. Yet, two primary component variables, leaf C:N ratio and sugar content varied along latitudinal and climatic gradients, respectively. The leaf C:N ratios were lowest at intermediate latitudes. This outcome may be due to temperature-related plant physiological stoichiometry and biogeographical gradients in soil substrate age (limitation of soil N at higher latitudes) (Reich & Oleksyn, 2004). Leaf soluble sugar content decreased with increasing precipitation (Cao *et al*., 2018). Soluble sugars, especially glucose and fructose, accumulate together with other osmolytes during drought (Nio *et al*., 2011), resulting in high concentration in areas where precipitation is low. The lack of variation of leaf defences contradicts the Latitudinal Herbivory Defence Hypothesis which predicts that plant species at lower latitudes experience higher mean rates of herbivory than their temperate counterparts (Coley & Barone, 1996; Schemske *et al*., 2009; Lim *et al*., 2015) and, for this reason, should have evolved higher levels of anti-herbivore defences (Rasmann & Agrawal, 2011; Pearse & Hipp, 2012). However, the generality of this hypothesis is currently under debate (Moles & Ollerton, 2016). Several studies found no evidence for a latitudinal gradient in herbivory and plant defences (Moles *et al*., 2011) while others did (Salgado & Pennings, 2005; Woods *et al*., 2012); there is also mixed evidence when comparing different herbivore species or plant defensive traits (Anstett *et al*., 2015; Moreira *et al*., 2015, 2018). A plausible explanation for the lack of latitudinal gradients in oak defences could be that we sampled leaves at the middle of the growing season rather than at the end, and we did not measure constitutive and induced defences separately. This is an insightful point because oak leaves may have differentially accumulated phenolics in response to herbivory (*i.e*., induced defences) or experienced marked differences in light intensity toward the end of the growing season (Karolewski *et al*., 2013). Furthermore, despite attempts to synchronize phenology across sites, seasonal changes in oak chemical defences (Salminen & Karonen, 2011) might have masked latitudinal patterns in defences. Therefore, further studies should include measurements at multiple time points during the growing season and distinguish between different types of defences, including physical vs. chemical defences (Wang *et al*., 2018) as well as constitutive vs. induced defences (Anstett *et al*., 2018) in order to address latitudinal gradients in plant defence more comprehensively.

We found no latitudinal or climatic gradients in bird attack rates on dummy caterpillars (**Figure 5**). These results agree with the large-scale study performed by Roslin *et al*. (2017) who found an increase of the activity of predatory arthropods in several plant species toward the equator, but no significant trend in avian predation. Several factors may explain the lack of response of avian predation to latitudinal or climatic gradients. First, bird species are distributed through migration allowing them to breed at higher latitudes, resulting in a constant predation rate across climatic and geographical clines (Dufour *et al*., 2020). In contrast, other predators with lower mobility such as arthropods (e.g. ants, ladybirds) are much more abundant at lower latitudes, resulting in a higher selection pressure toward the equator (Roslin *et al*., 2017). Second, bird communities are more influenced by forest habitat composition at lower latitudes, and more by food availability at higher latitudes (Charbonnier *et al*., 2016) where the diet variability is lower (Barnagaud *et al*., 2019), suggesting a stronger effect of local habitat features (e.g. resource availability and habitat suitability) than climatic gradients. Third, we cannot exclude that the lack of latitudinal trend in bird attack rates resulted from methodological limitations due to the fact we only exposed green dummy caterpillars, in low hanging branches. Birds depend more on food accessibility than abundance *per se*, so that the exact location of dummy caterpillars regarding factors such as edge, light contrast and shrubby understory may have modified the perception and the accessibility to the prey (Zvereva *et al*., 2019). Thus, the absence of a latitudinal trend in bird predation cannot be inferred only from geographical location.

### Mechanisms underlying latitudinal and climatic variation in incidence of gall-inducers and leaf-miners

The incidence of gall-inducers and leaf-miners was partially explained by variability in several leaf chemical traits (**Figure 5**). For instance, the incidence of gall-inducers increased with increasing leaf soluble sugars and N concentrations, which is consistent with gall-inducers being metabolic sinks (Huang *et al*., 2014). However, the effects of temperature and precipitation on leaf-miners were likely indirectly mediated by climatic variation in defences, as such effects became non-significant once condensed tannins and hydrolysable tannins were included in the model. These results agree with previous studies reporting indirect effects (via leaf defences) of climate on herbivory (Anstett *et al*., 2018; Moreira *et al*., 2018). For instance, Anstett *et al*. (2018) found indirect effects of climate on insect herbivory in 80 species of evening primroses, which were mediated by leaf chemicals (total phenolics and oenothein A). Contrarily, the effects of temperature and precipitation on gall-inducers were not indirectly mediated by climatic variation in defences, as in this case such effects remained significant after chemical traits were included in the models. In this sense, it is possible that other unmeasured defensive traits (e.g. physical defences), strategies (e.g. induced defences, tolerance) or indirect influence of parasitoids would have accounted for the observed climatic variation in the incidence of gall-inducers.

Our results showed that the effects of temperature and precipitation on incidence of gall-inducers and leaf-miners were not indirectly mediated by climatic variation in attack rate, as such effects remained significant after including bird attack rates in the models. This result is in line with Zverev *et al*. (2020) study performed in Arctic ecosystems, that concluded that birds are unlikely to shape the spatial patterns of insect herbivory. Associations between bird insectivory and insect herbivore can be positive (Mäntylä *et al*., 2014; Gunnarsson *et al*., 2018), negative (Maguire *et al*., 2015; Kozlov *et al*., 2017) or non-significant (Moreira *et al*., 2019; Valdés-Correcher *et al*., 2019), depending on the study and methods used. Arthropod predators (e.g. ants, ladybirds) play an important role on herbivore populations and may respond to large-scale variation in climatic conditions at greater extent than vertebrate predators (Roslin *et al*., 2017; Zvereva *et al*., 2019). For example, a meta-analysis conducted by Rodríguez-Castañeda (2013) found that ant predation on herbivores significantly increase at higher temperatures and precipitations, indicating that plants growing under warmer and wetter conditions exhibit lower levels of insect herbivory. Besides, birds are considered intraguild predators that not only eat insect herbivores but also arthropod predators (Gunnarsson, 2007) and intraguild predation may weaken herbivore suppression (Finke & Denno, 2005). Unfortunately, we were not able to quantify predation rates by such arthropods nor intraguild predation, which weakens our conclusions about the potential role of predators across climatic gradients.

### Conclusion

By simultaneously investigating bottom-up and top-down forces driving insect herbivory along latitudinal and climatic clines in a widespread tree species in Europe, this study brings new insights into the vivid debate about latitudinal variation in the direction and strength of biotic interactions (Schemske *et al*., 2009; Moles *et al*., 2013; Anstett *et al*., 2016; Roslin *et al*., 2017). We found no evidence that latitude or climate influenced leaf-chewing herbivores, but we found that climatic factors rather than latitude *per se* were the best predictors of the large-scale variation in the incidence of leaf-miner and gall-inducer herbivores as well as in variation in leaf nutritional quality. In sharp contrast, we found no evidence that plant chemical defences and bird attack rates were influenced by latitude or climatic factors, which conflicts with the dominant view in ecology (Moles & Ollerton, 2016; Roslin *et al*., 2017; Zvereva *et al*., 2019). Because unravelling causes of latitudinal variation in the strength of biological interactions is one of the common approaches for the prediction of biotic interactions under global warming (Verheyen *et al*., 2019), it is crucial that future studies simultaneously test for effects of latitude *per se* and climate on insect herbivory by different feeding guilds (Kozlov *et al*., 2017), as well as investigate the mechanisms underlying such effects.

## Acknowledgements

This study has been carried out with financial support from the French National Research Agency (ANR) in the frame of the Investments for the future Programme, within the Cluster of Excellence COTE (ANR-10-LABX-45). The authors warmly thank all young European citizens and their teachers who have made this study possible. They also thank professional scientists who have kindly accepted to participate in this study: Stefan K. Müller (Freie evangelische Schule Lörrach), Olga Mijón Pedreira (teacher IES Rosais 2, Vigo-Spain) and Mickael Pihain (Research Unit “Ecosystèmes, Biodiversité, Evolution”, University of Rennes 1 / CNRS, 35042 Rennes, France), and Chloe Mendiondo and Claire Colliaux (Department of Agroecology, Aarhus University, Flakkebjerg Research Centre, DK-4200 Slagelse, Denmark). The authors declare no competing financial interests.

