## Supplementary material for "Search for top-down and bottom-up drivers of latitudinal trends in insect herbivory in oak trees in Europe"

**Supplementary material S1**

Detailed methodology

***Leaf phenolics* -** We quantified leaf phenolics only on leaves collected by professional scientists in 2018. For each tree, we selected 10 mature leaves with no evidence of insect damage and we grounded them to thin powder. Then, we extracted phenolic compounds from 20 mg of dry plant tissue with 1 mL of 70% methanol in an ultrasonic bath for 15 min. We centrifuged and subsequently transferred them to chromatographic vials. To perform the chromatographic analyses we used Ultra-High-Performance Liquid-Chromatograph (UHPLC Nexera LC-30AD; Shimadzu) equipped with a Nexera SIL-30AC injector and one SPD-M20A UV/VIS photodiode array detector. The compound separation was carried out on a Kinetex™ 2.6 µm C18 82-102 Å, LC Column 100 × 4.6 mm, protected with a C18 guard cartridge. The flow rate was 0.4 mL min-1 and the oven temperature was set at 25 °C. The mobile phase consisted of two solvents: water-formic acid (0.05%) (A) and acetonitrile-formic acid (0.05%) (B), starting with 5% B and using a gradient to obtain 30% B at 4 min, 60% B at 10 min, 80% B at 13 min and 100 % B at 15 min. The injection volume was between 15-30 µL (from a total of 24 samples we injected 30 µL because the concentration of secondary metabolites was quite low).

We identified four groups of phenolic compounds: flavonoids, ellagitannins and gallic acid derivates (“hydrolysable tannins” hereafter), proanthocyanidins (“condensed tannins” hereafter) and hydroxycinnamic acid precursors to lignins (“lignins” hereafter).We quantified flavonoids as rutin equivalents, condensed tannins as catechin equivalents, hydrolysable tannins as gallic acid equivalents, and lignins as ferulic acid equivalents (Moreira *et al.*, 2018). We obtained the quantification of these phenolic compounds by external calibration using calibration curves at 0.25, 0.5, 1, 2 and 5 μg mL-1. Phenolic compound concentrations were expressed in mg·g-1 tissue on a dry weight basis.

***Nutritional traits* -** We quantified plant nutritional quality on leaves collected by both professional scientists and schoolchildren. We grounded the 60 oven dried leaves on which we scored herbivory to thin powder such that leaf nutritional traits reflected the content of leaves with different amounts of herbivore damage.

We quantified macroelements (C, N, P) after wet mineralisation (H2SO4+H2O2). Phosphorus and nitrogen were quantified colorimetrically with an AutoAnalyser 3 High Resolution colorimeter (SEAL), using ammonium molybdate (for P) and sodium salicylate (for N) as reagents. We also quantified leaf C:N ratio with a gas chromatography in an automatic elemental analyser (FlashEA 1112; Thermo Fisher Scientific Inc.) using 6 µg of dried leaf powder.

We purified between 0.1 and 0.5 g of dried leaf powder to holocellulose using the Jayme–Wise method (Leavitt & Danzer, 1993). Leaf powder was placed in a Teflon bag and sequentially treated in a Soxhlet extractor with 2:1 toluene:ethanol, then 100% ethanol, to remove extractables. The samples were then boiled in water to remove soluble carbohydrates, and bleached at a temperature of 70°C in 4 mL of acetic acid solution with 21 g of sodium chlorite to decompose the lignin. The samples were weighed and this weight corresponded with the content on cellulose.

We extracted soluble sugars from 50 mg of dried leaf powder. The dry material was transferred to a tube (tube A) with 1 mL of ethanol in a water bath for 30 min at 80°C. We centrifuged and subsequently transferred the liquid to an Eppendorf (tube B). We added 1 mL of 50% ethanol in the tube A and placed it in water bath for 30 min at 80°C. We centrifuged again and subsequently transferred the liquid to the tube B. We added 1 mL of 20% ethanol in the tube A and placed it in water bath for 30 min at 80°C. We centrifuged and subsequently transferred the liquid to the tube B. We added 1 mL of NaOH 0.02N in the tube A and placed it in water bath for 30 min at 90°C. We centrifuged and subsequently transferred the liquid from the tube B to the tube A. Both tubes were placed in a speed vac for complete evaporation. Then, 50 µL aliquots of the diluted solutions were injected into 2.5mL of anthrone reagent which allows colorimetric analysis of the total sugar content (all monosaccharides, disaccharides and polysaccharides in their hydrolysed or non-hydrolysed forms). The preparation of the anthrone reagent was adapted from Bachelier and Gavinelli (1966): 0.5 g of anthrone was directly dissolved in 250mL of sulphuric acid at 95–98%. The colorimetric reaction was accelerated by heating at 80°C for 30 min and the total sugar content was then determined by measuring the absorbance at 560nm with a spectrophotometer (Biochrom Libra S22, Biochrom, Cambridge, UK). The sugar concentration was determined from calibration curves established using standard sucrose solutions with a range of known concentrations.

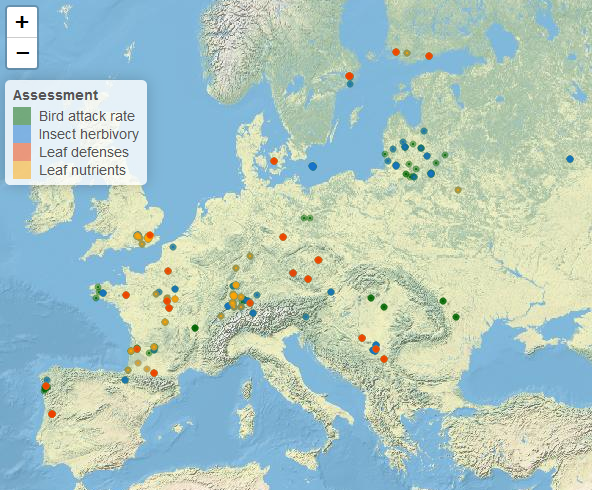

**Figure S1**. Location of trees sampled for the assessment of herbivory, predation attack rate, leaf nutrients and defences.

Interactive version of the maps:

**Figure S1A.** Trees sampled for the assessment of herbivory.

**Figure S1B.** Trees sampled for the assessment of predation attack rate.

**Figure S1C.** Trees sampled for the assessment of leaf nutrients.

**Figure S1D.**Trees sampled for the assessment of leaf defences.

**Figure S1E.** Trees where the assessment of herbivory, predation attack rate, lead nutrients and leaf defences was assessed simultaneously.

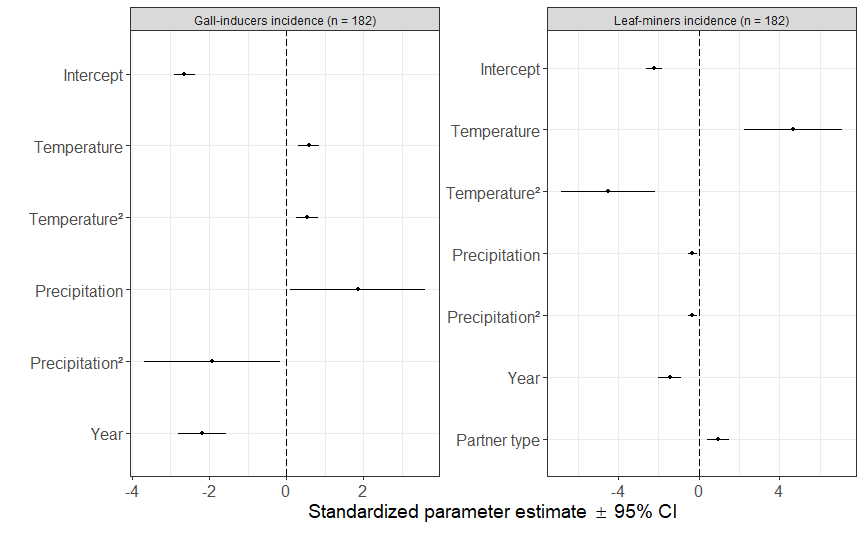

**Figure S2.** Effects of partner type, year, temperature and precipitation on gall-inducers and leaf-miners incidences. Circles and error bars represent standardized parameter estimates and corresponding 95% CI. The vertical dashed line centered on zero represents the null hypothesis. Small and bigl circles indicate significant and non-significant effect sizes, respectively.

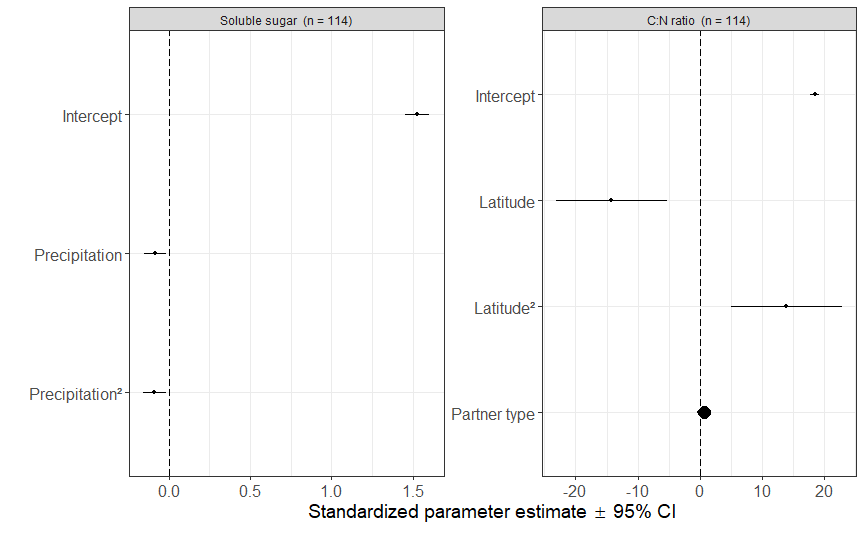

**Figure S3**. Effects of partner type, year, latitude, longitude, temperature and precipitation on leaf C:N ratio and leaf soluble sugar (g L⁻¹). Circles and error bars represent standardized parameter estimates and corresponding 95% CI. The vertical dashed line centered on zero represents the null hypothesis. Small and big circles indicate significant and non-significant effect sizes, respectively.

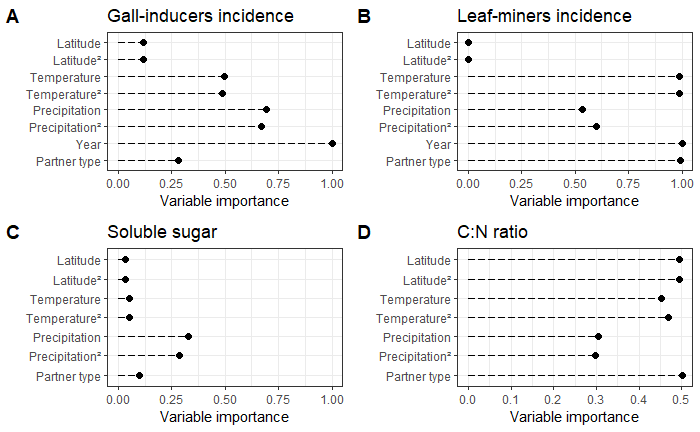

**Figure S4**. Variable importance of every variable included in the geographic and climatic models that considered the effect of longitude, latitude, temperature and precipitation on herbivory (gall-inducers and leaf-miners incidences) and on leaf chemistry.

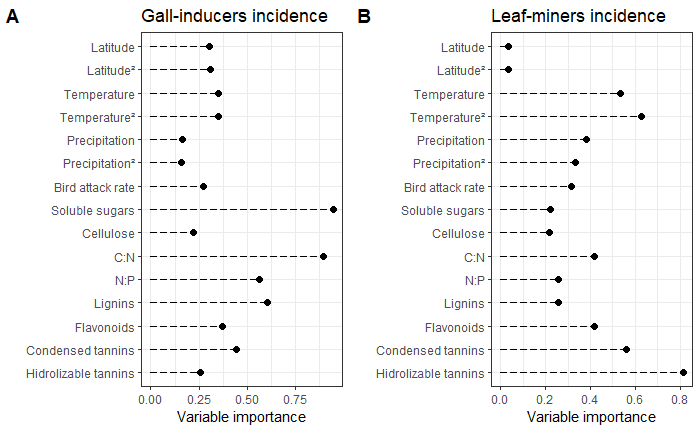

**Figure S5**. Importance of every variable included in the biotic model that considered the effect of leaf traits, bird attack rate, climatic variables on gall-inducers and leaf-miners incidence.

**Table S1**. Summary of the different variables measured.

| Variables | Mean (n, sd) | |
| --- | --- | --- |
|  | Scientific partner | School partner |
| Tree height (m) | 14.75 (97, 7.06) | 13.01 (126, 7.45) |
| Tree circumference (cm) | 121.35 (97, 79.81) | 103.94 (126, 93.71) |
| Herbivory (%) | 9.55 (104, 6.64) | 7.46 (78, 4.33) |
| Gall-inducers incidence | 0.08 (104, 0.09) | 0.05 (78, 0.09) |
| Leaf-miners incidence | 0.24 (104, 0.19) | 0.10 (78, 0.11) |
| Bird attack rate | 0.02 (115, 0.01) | 0.01 (137, 0.01) |
| Soluble sugar (g L⁻¹) | 3.51 (72, 1.49) | 4.09 (42, 2.09) |
| Cellulose (g) | 0.09 (72, 0.04) | 0.12 (42, 0.05) |
| C:N ratio | 19.0 (72, 2.56) | 18.04 (42, 2.17) |
| N:P ratio | 17.22 (72, 5.55) | 14.82 (42, 2.88) |
| Lignin (mg g⁻¹ ) | 1.05 (78, 1.23) |  |
| Hydrolysable tannins (mg g⁻¹ ) | 0.47 (78, 0.54) |  |
| Condensed tannins (mg g⁻¹ ) | 1.25 (78, 1.08) |  |
| Flavonoids (mg g⁻¹ ) | 2.12 (78, 2.07) |  |
| Total defences (mg g⁻¹ ) | 4.89 (78, 4.30) |  |

[**Table S**](https://drive.google.com/file/d/1KoB18iknBoJ5CHZGD6h5ivFHvB-Q_jAQ/view?usp=sharing)**2**. We included model parameters loglink, AICc, delta and weight of the different climatic models.

[**Table S**](https://drive.google.com/file/d/1cs79am21QX1AjqdNepvf73TU2zIHceY4/view?usp=sharing)**3.** We included model parameters  loglink, AICc, delta and weight of the different biotic models.
